## Supplementary Data 1 for "Identification of a RAB32-LRMDA-Commander membrane trafficking complex reveals the molecular mechanism of human oculocutaneous albinism type 7"

Raw blots for Figure 1

1D

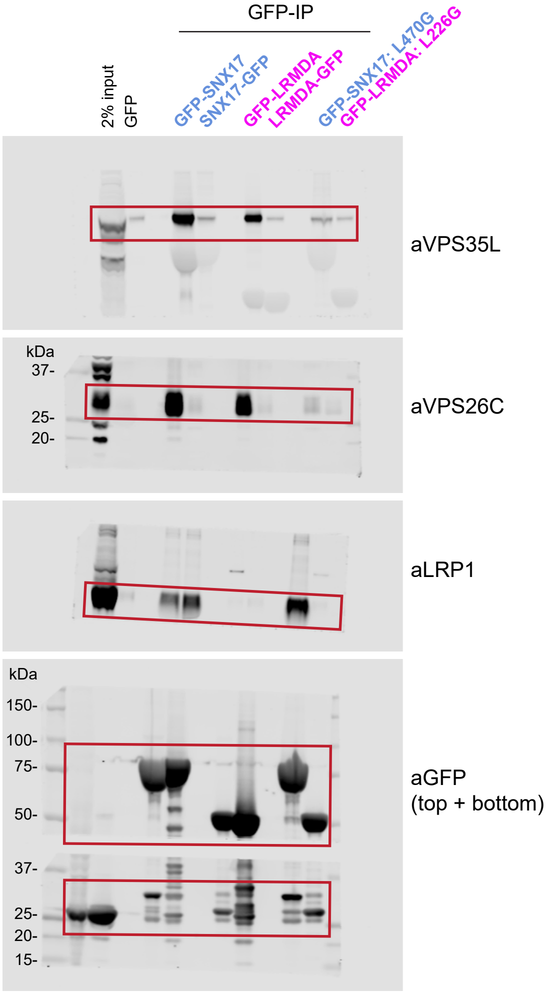

### Raw blots for Figure 2

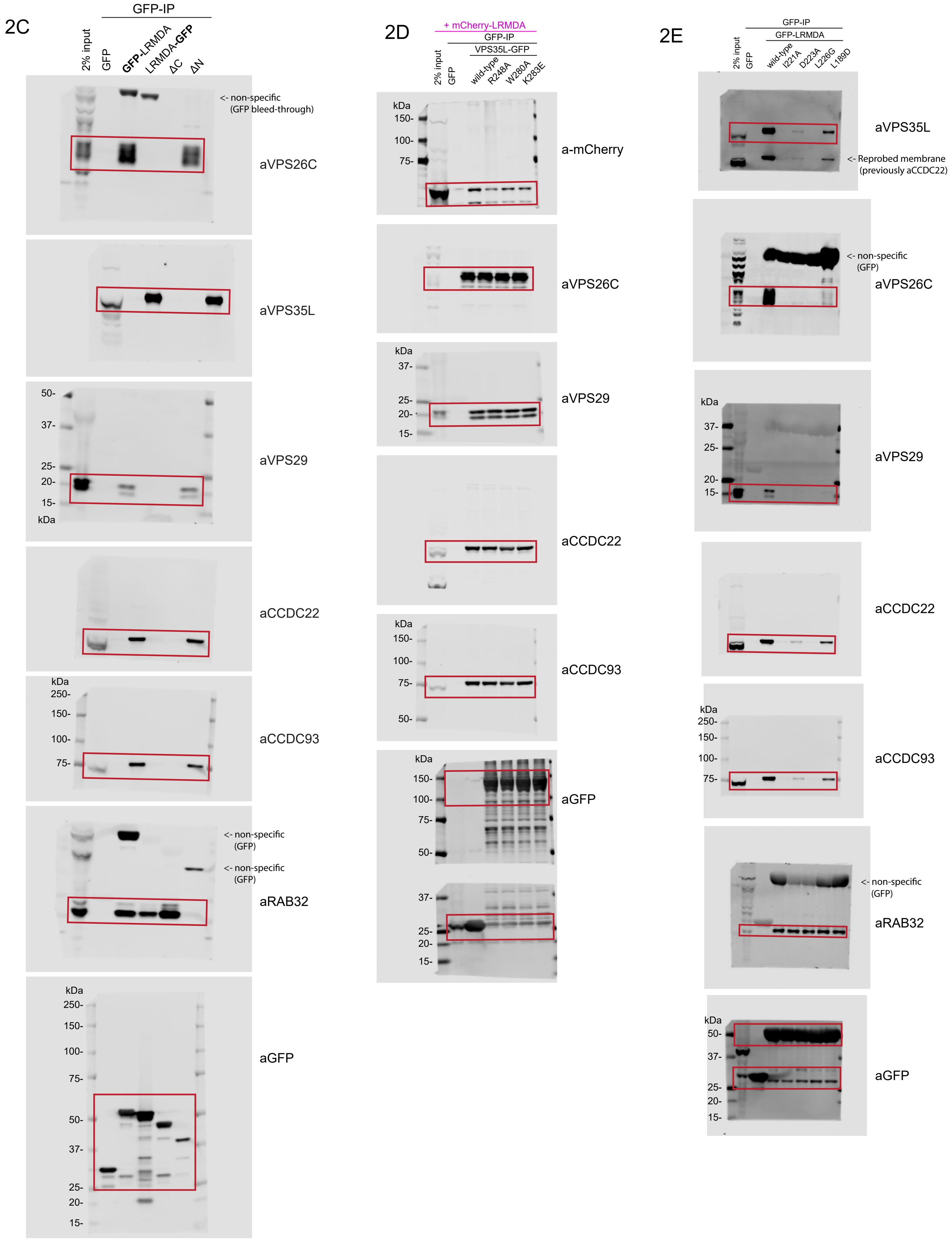

### Raw blots for Figure 3

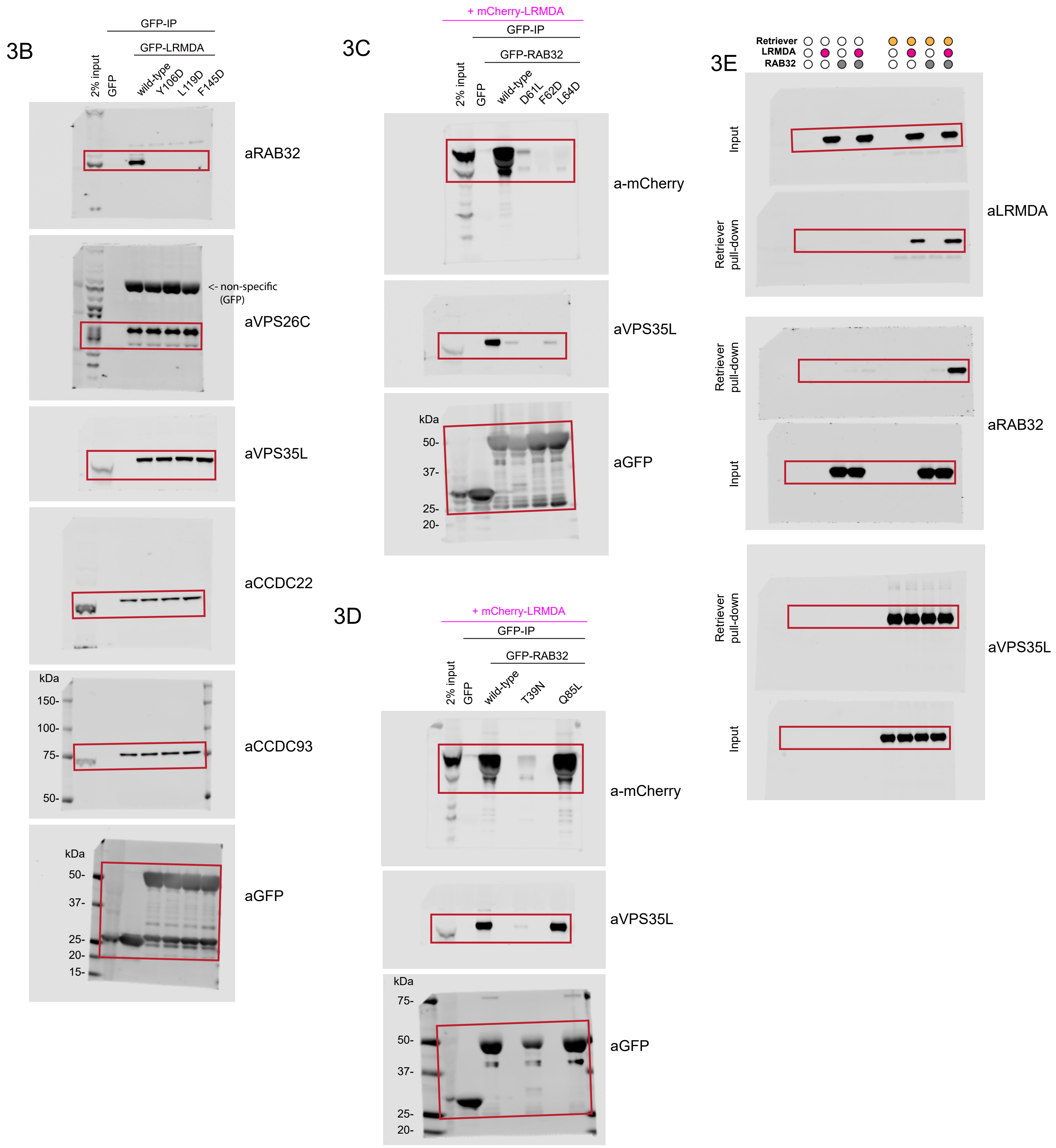

Raw blots for Figure 4

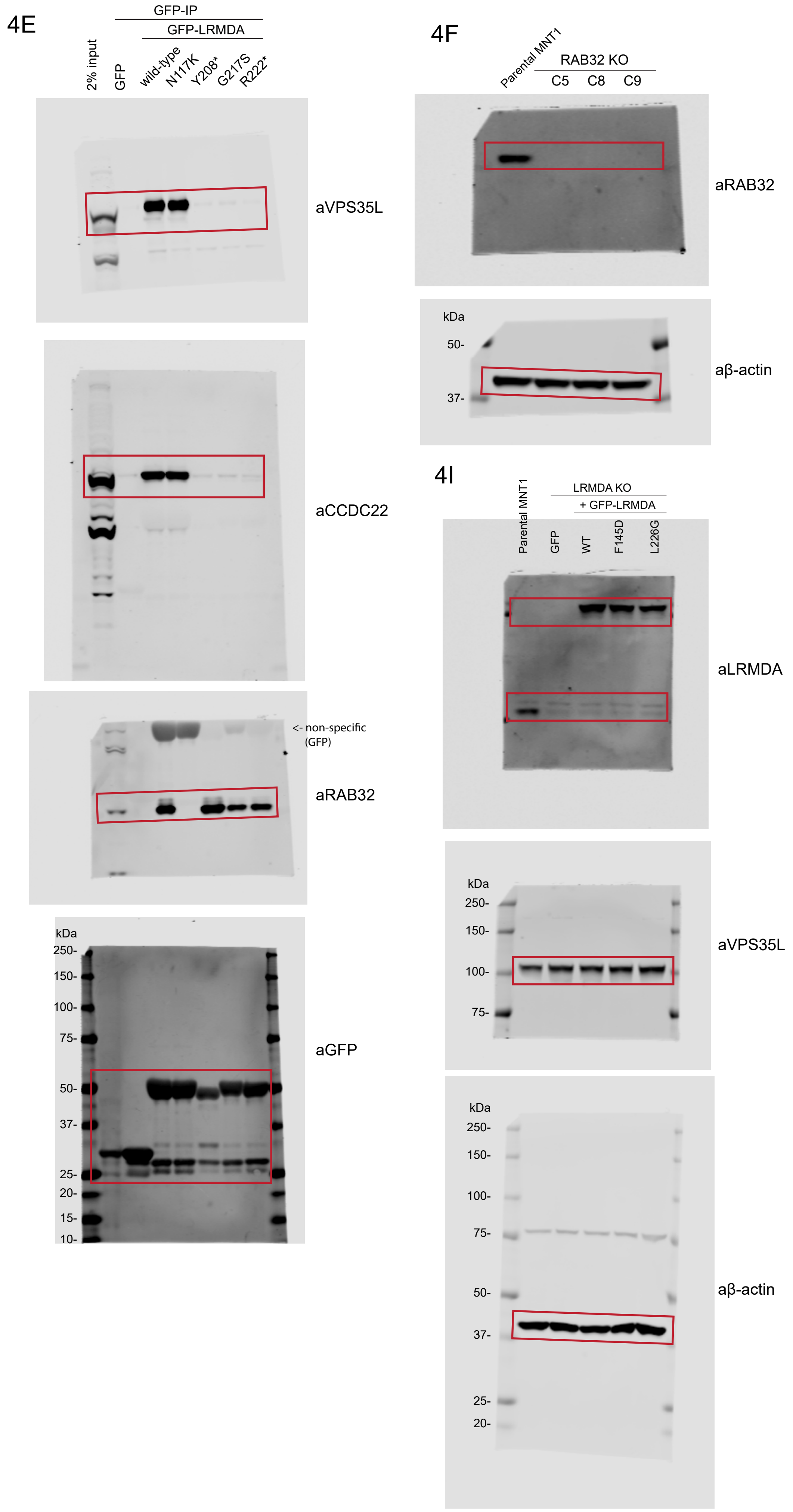

### Raw blots for Figure 5

## 5A

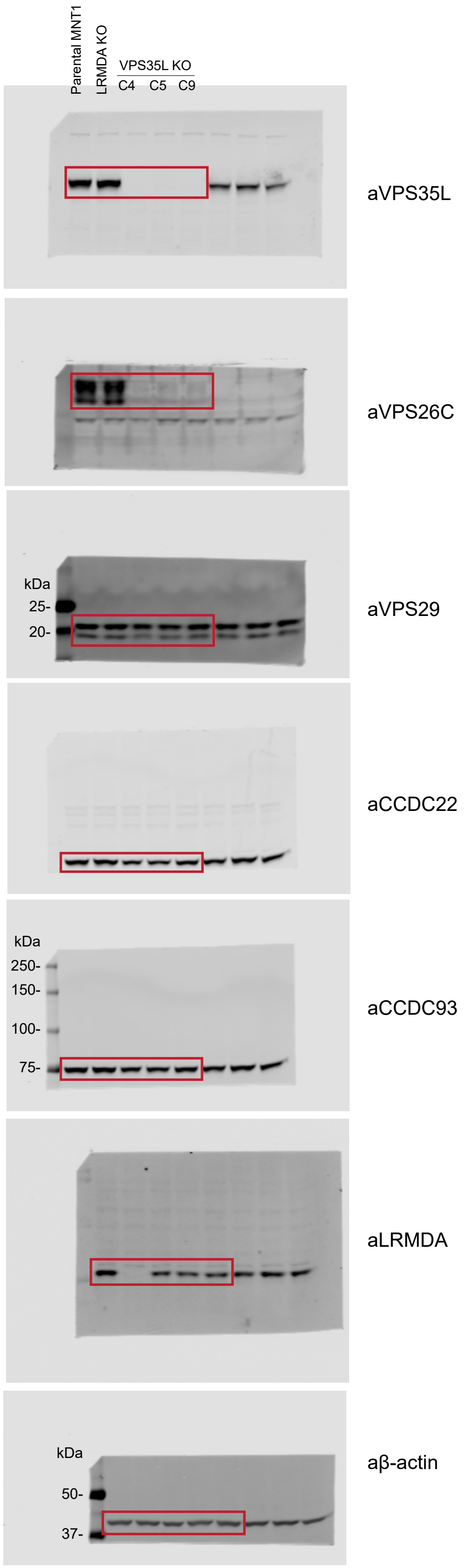

Raw blots for Figure 6

6B

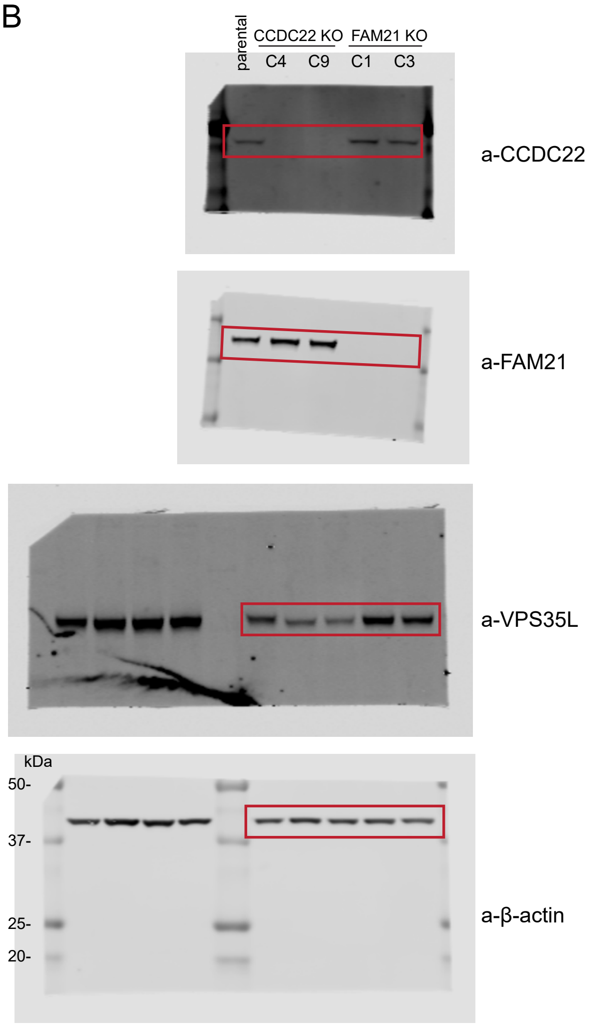


6E

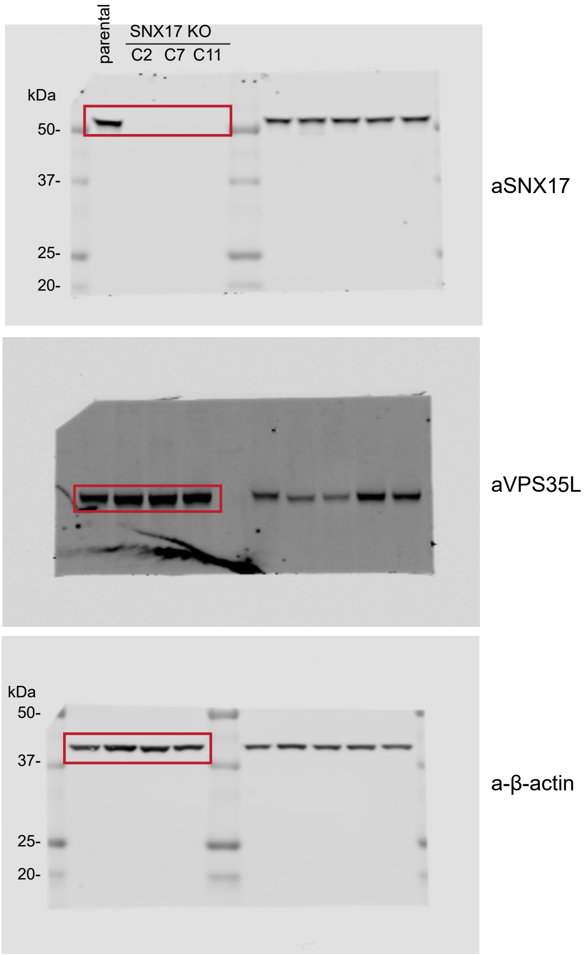

### Raw blots for Figure 7

## 7B

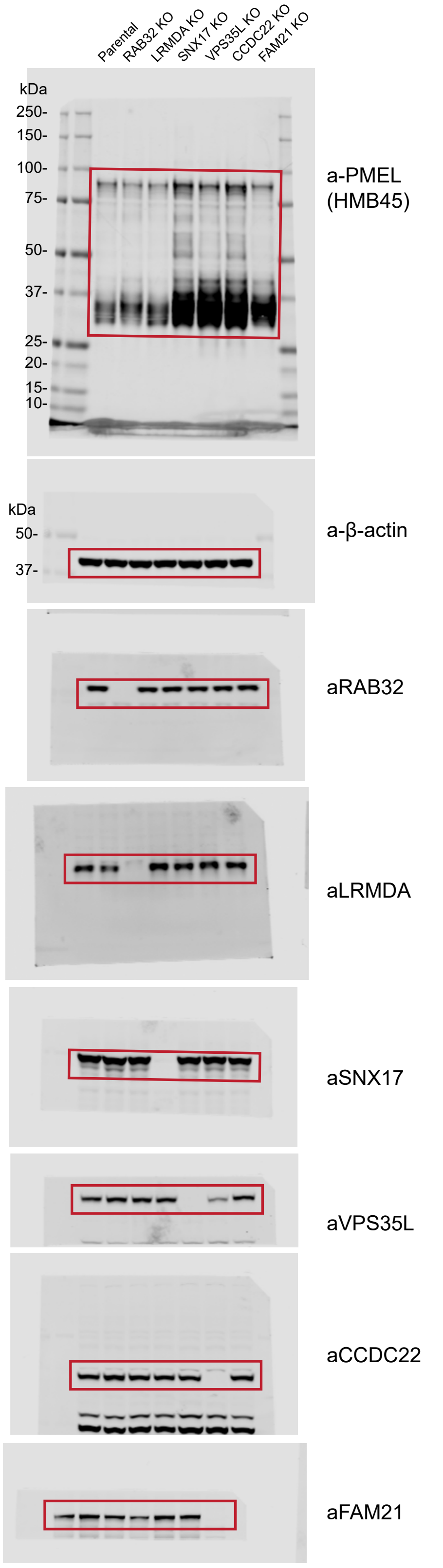

### Raw blots for Supplementary Figure 2

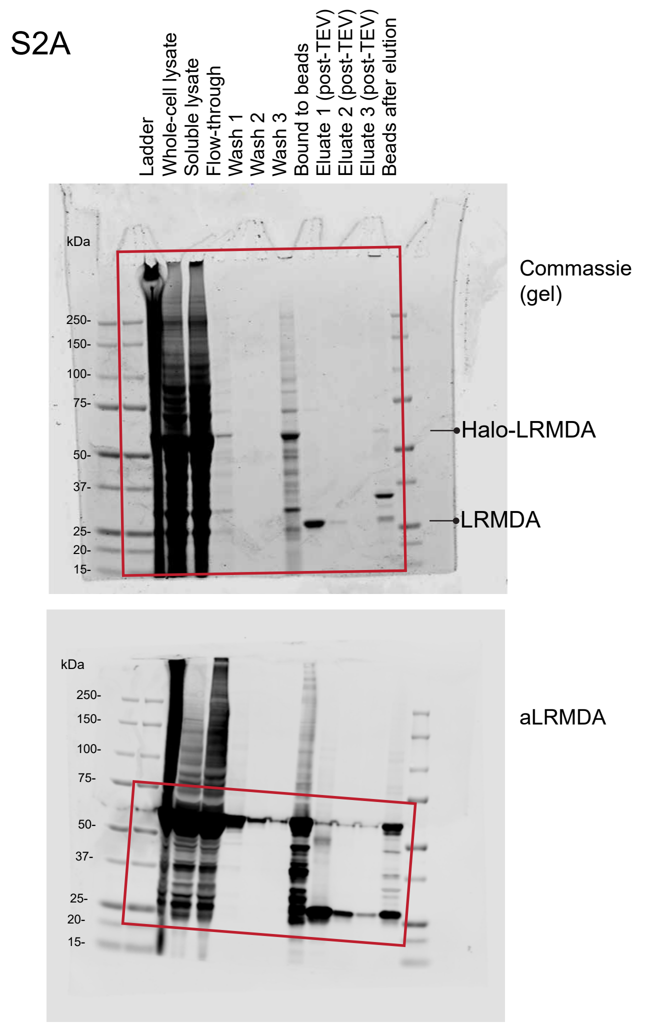


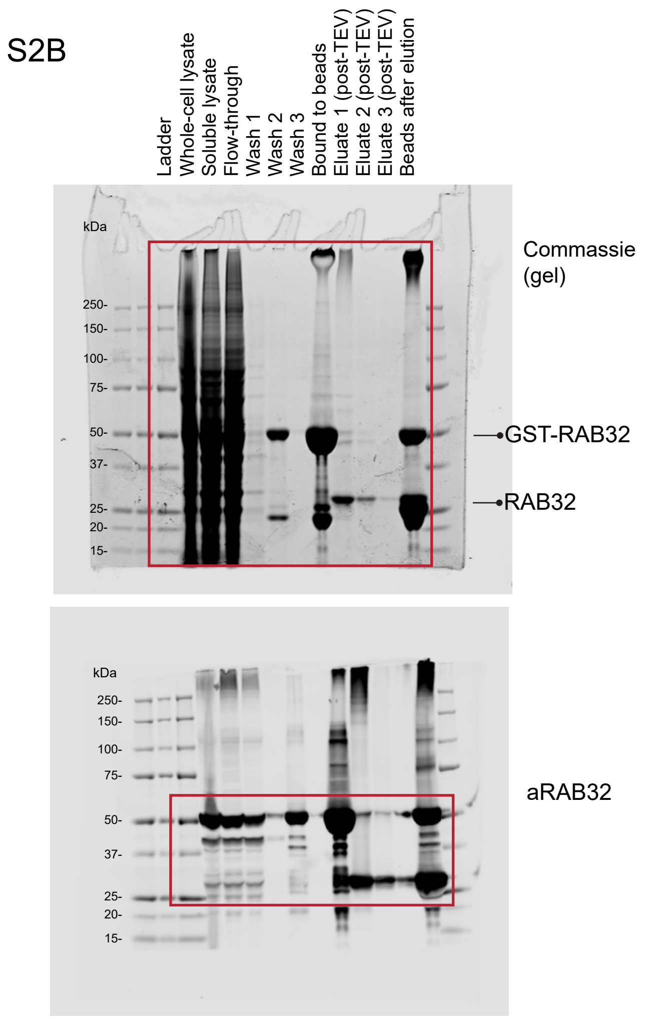


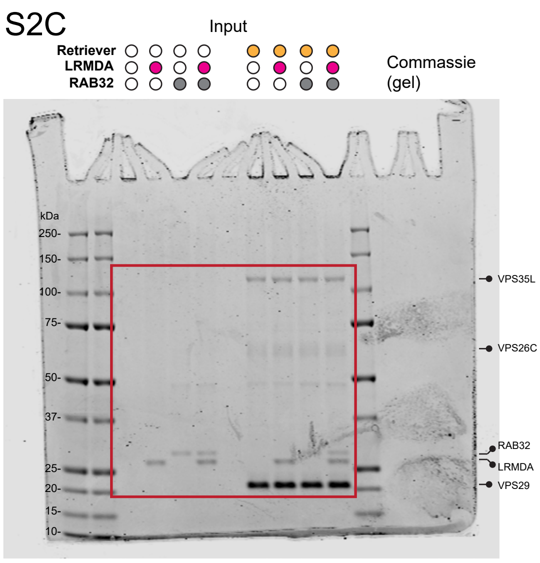


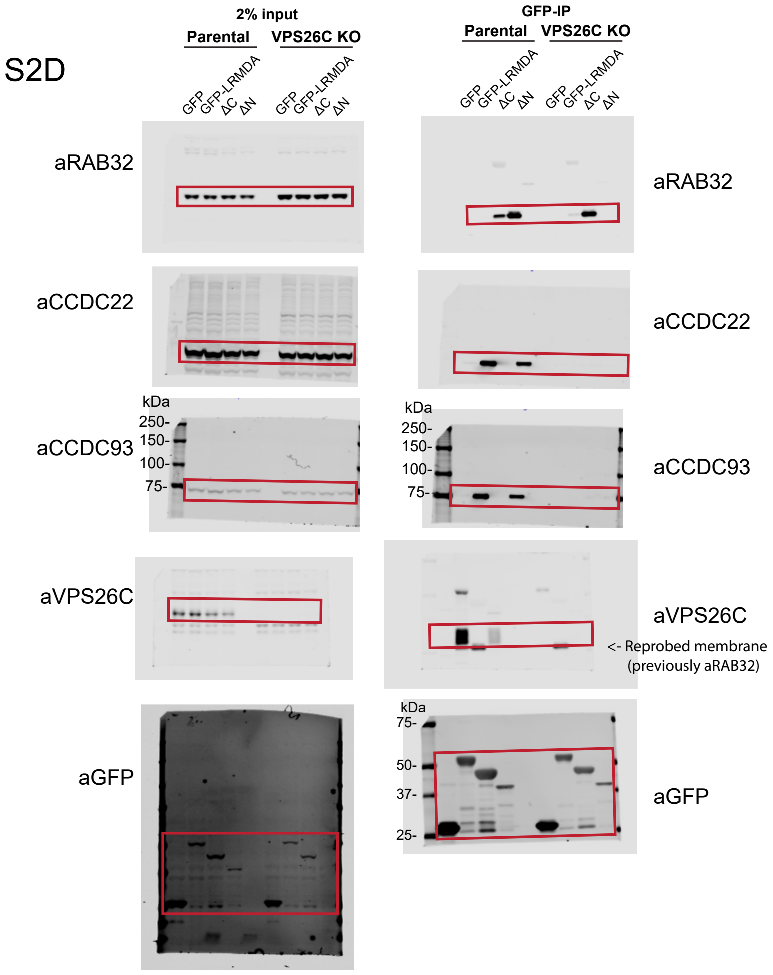

### Raw blots for Supplementary Figure 3

## S3B

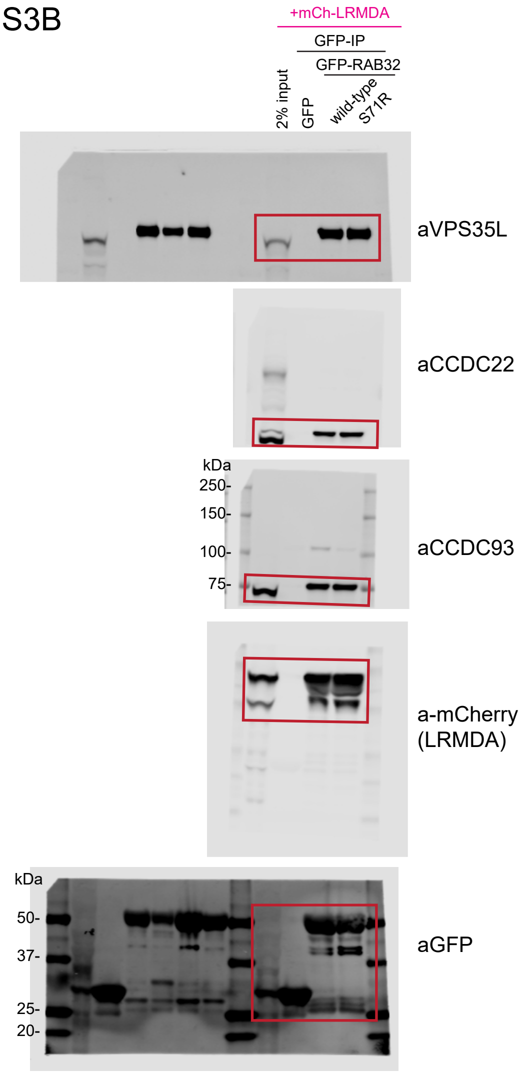


## S3D

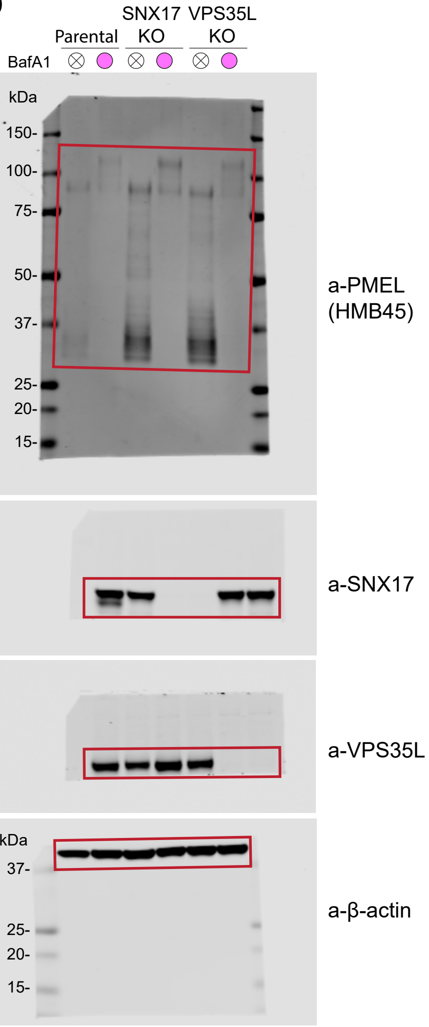
